# The landscape of somatic mutation in normal colorectal epithelial cells

**DOI:** 10.1101/416800

**Authors:** Henry Lee-Six, Peter Ellis, Robert J. Osborne, Mathijs A. Sanders, Luiza Moore, Nikitas Georgakopoulos, Franco Torrente, Ayesha Noorani, Martin Goddard, Philip Robinson, Tim H. H. Coorens, Laura O’Neill, Christopher Alder, Jingwei Wang, Rebecca C. Fitzgerald, Matthias Zilbauer, Nicholas Coleman, Kourosh Saeb-Parsy, Inigo Martincorena, Peter J. Campbell, Michael R. Stratton

**Affiliations:** Wellcome Trust Sanger Institute, Hinxton, UK; Department of Hematology, Erasmus University Medical Center, Rotterdam, The Netherlands; Department of Surgery and Cambridge NIHR Biomedical Research Centre, Cambridge Biomedical Campus, Cambridge, UK; Department of Paediatric Gastroenterology, Cambridge University Hospital Trust, Addenbrookes, Cambridge, UK; Medical Research Council Cancer Unit, Hutchison/Medical Research Council Research Centre, University of Cambridge, Cambridge, UK, and Cambridge University Hospitals NHS Trust, Hills Road, Cambridge, UK; Department of Pathology, Papworth Hospital NHS Trust, UK; Department of Pathology, University of Cambridge, Cambridge, UK and Cambridge University Hospitals NHS Foundation Trust, Cambridge, UK

## Abstract

The colorectal adenoma-carcinoma sequence has provided a paradigmatic framework for understanding the successive somatic genetic changes and consequent clonal expansions leading to cancer. As for most cancer types, however, understanding of the earliest phases of colorectal neoplastic change, which may occur in morphologically normal tissue, is comparatively limited because of the difficulty of detecting somatic mutations in normal cells. Each colorectal crypt is a small clone of cells derived from a single recently-existing stem cell. Here, we whole genome sequenced hundreds of normal crypts from 42 individuals. Signatures of multiple mutational processes were revealed, some ubiquitous and continuous, others only found in some individuals, in some crypts or during some phases of the cell lineage from zygote to adult cell. Likely driver mutations were present in ∼1% of normal colorectal crypts in middle-aged individuals, indicating that adenomas and carcinomas are rare outcomes of a pervasive process of neoplastic change across morphologically normal colorectal epithelium.

## Introduction

Sequencing of >20,000 cancers has identified the repertoire of driver mutations in cancer genes converting normal cells into cancer cells and revealed the mutational signatures of the underlying biological processes generating somatic mutations^1,2^. Cancers are, however, end stages of an evolutionary process operating within cell populations and commonly arise through the accumulation of multiple driver mutations engendering a series of clonal expansions. Understanding this progression has depended, in substantial part, on identifying somatic mutations in morphologically abnormal neoplastic proliferations representing intermediate stages between normal and cancer cells. Classical studies of driver mutations in colorectal adenomas and carcinomas have been particularly influential in shaping our perspective in this regard^3^.

As for most cancer types, however, the earliest stages of progression to colorectal cancer remain considerably less well understood. The driver mutation that first sets a colorectal epithelial cell on the path to cancer is likely caused by mutational processes operative in normal cells, of which there is limited understanding. The nature and numbers of the earliest neoplastic clones with driver mutations, which conceivably are morphologically indistinguishable from normal cells, are similarly unclear. In large part, these deficiencies are due to the technical challenge of identifying somatic mutations in normal tissues, which are composed of myriad microscopic cell clones. Several approaches have been adopted to address this, including sequencing of *in vitro* expanded cell populations derived from single cells^4-9^, sequencing normal tissue microbiopsies incorporating small numbers of clones^10,11^, sequencing single normal cells^12-14^, highly error corrected sequencing^15^, and non-sequencing based approaches^27,44^.

These approaches have provided insights into early stages of cancer development. Signatures of common somatic mutational processes have been found in normal cells of the small and large intestine, liver, blood, skin, and nervous system but thus far studies have not been of sufficient scale to characterise variation in their activity or detect less frequent processes^4-10,12-15.^ Remarkably high proportions of normal skin epithelial cells have been shown to be members of clones already carrying driver mutations^10^, and large mutant clones have been detected in blood^16-19^. Driver mutations have similarly been detected in a high proportion of endometrial crypts^11^. The extent of this phenomenon in the colon, an organ with a high cancer incidence, has not been investigated.

The colonic epithelial lining is a contiguous cell sheet organised into ∼15,000,000 glandular units, known as crypts, oriented perpendicular to the luminal surface and composed of ∼2,000 cells^20^. Towards the base of each crypt resides a small number of stem cells ancestral to the maturing and differentiated cells in the crypt^21^. These stem cells stochastically replace one another through a process of neutral drift^22,23^ such that all stem cells, and thus all cells, in a crypt derive from a single ancestor stem cell that existed in recent years^24-27^. The somatic mutations that were present in this ancestor are thus found in all ∼2,000 descendant cells and can be revealed by DNA sequencing of an individual crypt. Following acquisition of the requisite numbers and combinations of driver mutations, these stem cells are also thought to be the cells of origin of colorectal cancers^28^. To characterise the earliest stages of colorectal carcinogenesis, somatic mutation burdens, mutational signatures and the frequency of driver mutations in normal colorectal epithelium were explored by sequencing individual colorectal crypts.

## Results

### Somatic mutations and mutational signatures

2,035 individual colonic crypts from 42 individuals aged 11 to 78 were isolated using laser capture microdissection and sequenced using a modified library-making protocol developed for small amounts of input DNA. The samples were from seven transplant organ donors, 34 individuals biopsied to investigate potential colorectal disease and an autopsy of a subject with oesophageal cancer. In total, 15 had colorectal cancer and 27 showed no evidence of colorectal disease (Supplementary Table). Samples from all individuals in this study are referred to as “normal” crypts as when a cancer was present only biopsies distant from the lesion were used. The distribution of the variant allele fractions of mutations from whole genome sequencing of 571 individual crypts indicated that the large majority of crypts were predominantly clonal cell populations derived from a single ancestral stem cell (Extended Data Fig. 1d). There was substantial variation in mutation burdens between individual crypts,ranging from 1,508 to 15,329 for individuals in their sixties, which was not obviously attributable to technical factors. To explore the biological basis of this variation we extracted mutational signatures and estimated the contribution of each to the mutation burden of each crypt (Methods, Supplementary Results).

Nine single base substitution (SBS), six doublet base substitution (DBS), and five small indel (ID) mutational signatures were found. Of these, 14 closely matched (Methods) a known reference signature (SBS1, SBS2, SBS5, SBS13, SBS18, DBS2, DBS4, DBS6, DBS8,DBS9, DBS11, ID1, ID2, and ID5, nomenclature as in Alexandrov et al^1^) and six did not (SBSA, SBSB, SBSC, SBSD, IDA, and IDB) (Fig. 1, Extended Data Fig. 2-4). Thus, new mutational signatures were extracted despite extensive prior analysis of cancers, perhaps due to masking by the comparative complexity of signature mixtures present in cancer genomes.

**Figure 1.**
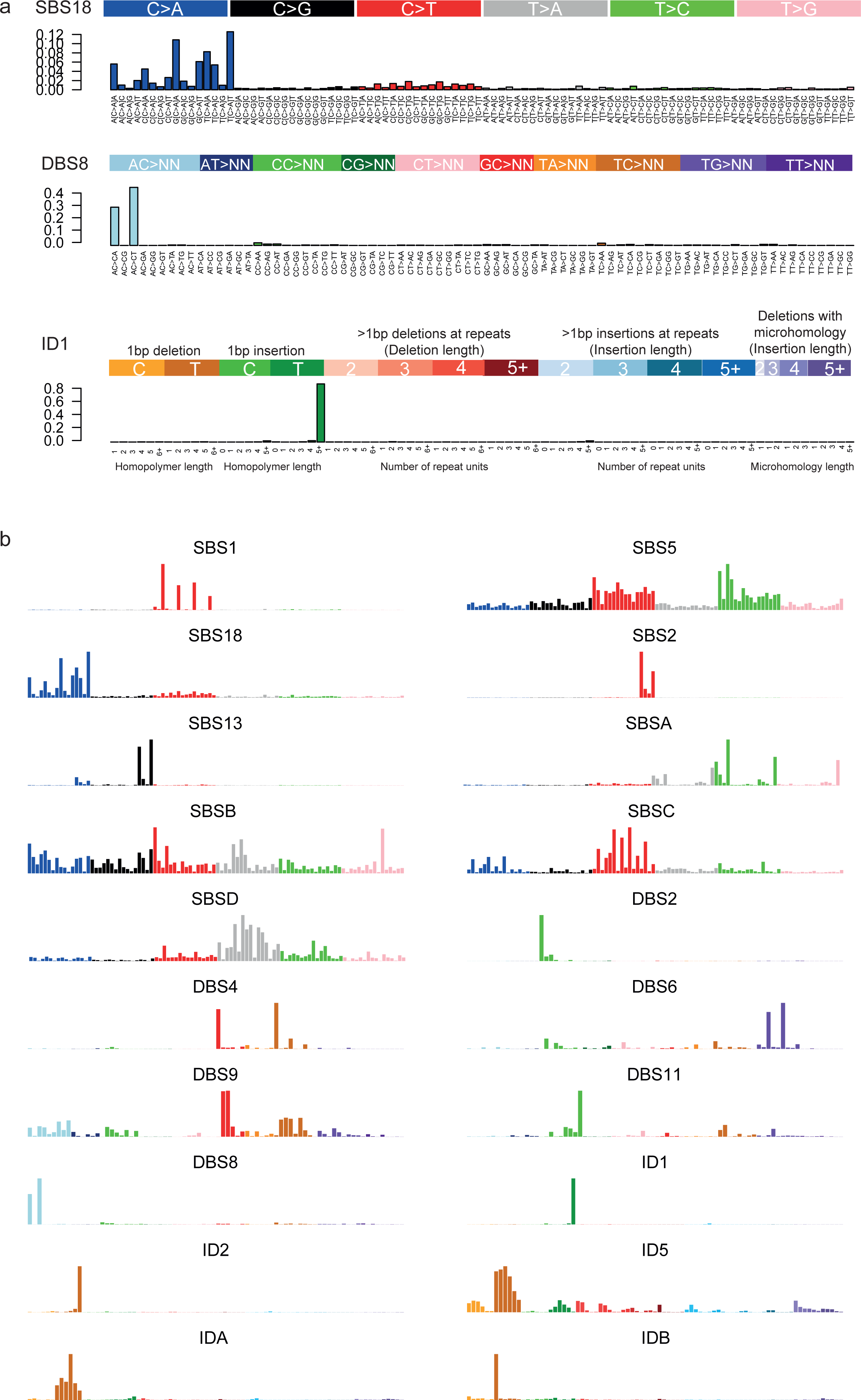
Mutational signatures present in normal colon. **a**, an example SBS, DBS, and ID signature showing the categories into which mutations are divided. Later figures are shown with the same categories, ordering, and colour scheme. **b**, the complement of signatures discovered in normal colonic epithelium. Known signatures are labelled according to their nomenclature in PCAWG, while novel signatures are labelled with letters. SBS, single base substitution; DBS, doublet base substitution; ID, small insertion or deletion.

### Ubiquitous mutational signatures

11 signatures (three SBS, five DBS and three ID) were found in >85% of crypts and are here termed “ubiquitous”. All have been previously described^1^. SBS1 is characterised by C>T substitutions at N<u>C</u>G trinucleotides (the mutated base is underlined) and is likely due to deamination of 5-methylcytosine. Its mutation load correlated linearly with age (Fig. 2). There was, however, variation in SBS1 mutation burdens between crypts from the same individual. This was due, in part, to different SBS1 mutation rates in different colonic sectors, with 16.8 mutations per year (95% CI 15.2-18.3) in the right (ascending and caecum), 16.1 (95% CI 14.4-17.5) in the transverse, and 12.8 (95% CI 11.1-14.4) in the left colon (descending and sigmoid). The SBS1 mutation rate in the terminal part of the small bowel, the ileum, was 12.7 (95% CI 10.6-14.9) (Supplementary Results).

**Figure 2.**
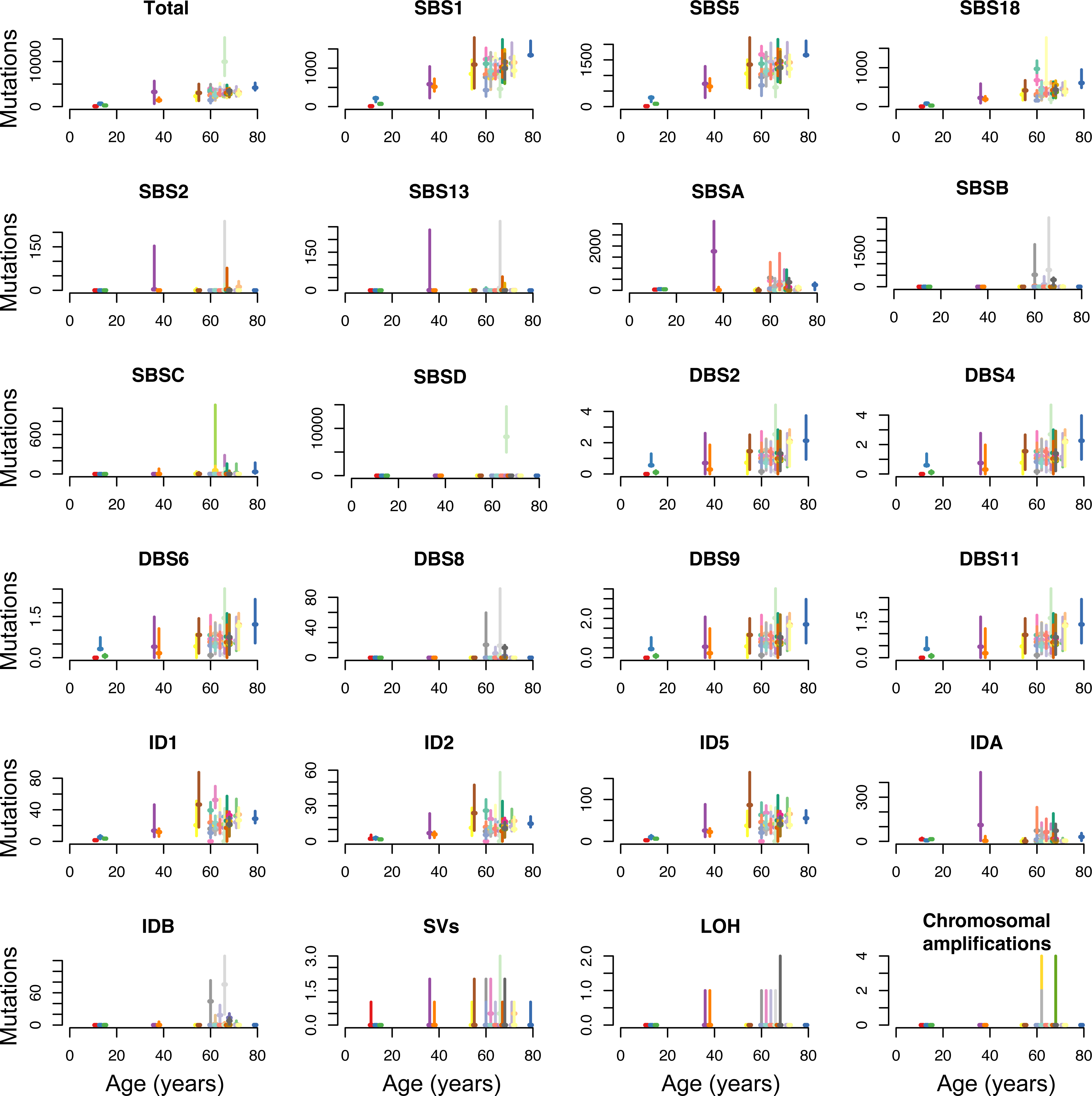
Mutation burden *versus* age for every signature. For every signature, the median (horizontal bar) and range (vertical bar) in mutation burden for all the crypts from each individual are shown. Each individual is coloured differently. See Supplementary Results for plots showing every crypt.

SBS5 is a relatively flat, featureless signature of unknown cause and SBS18 is predominantly characterised by C>A mutations, which may be due to DNA damage by reactive oxygen species^29,30^. The mutation burdens of these signatures also showed positive correlations with age, with the same ordering of sector differences as SBS1. Even after taking anatomical location and age into account, differences in mutation burden remained between different crypts, notably for SBS18, indicating that additional factors influence mutation rates in normal cells (Fig. 2, Extended Data Fig. 9, Extended Data Fig. 6al).

DBS2, DBS4, DBS6, DBS9, and DBS11 were tightly correlated in all colonic crypts. A composite spectrum of DBS2 and DBS4 is also present in normal mouse cells and, in human cancers, both correlate with age of diagnosis confirming that a substantial proportion of their mutations are generated in normal cells^1^. ID1, ID2, and ID5, which are predominantly characterised by insertions and deletions of a single T and may be the consequence of slippage during DNA replication, all accumulated linearly with age with the same order of sector differences as SBS1, SBS5, and SBS18 (Supplementary Results, Extended Data Fig. 5).

The correlations of mutation burden with age indicate that the mutational processes underlying these ubiquitous mutational signatures operate continuously throughout life, in all individuals and in all colorectal stem cells at similar rates. However, the results also suggest that differences in physiology and/or microenvironment (and potentially age of the most recent common ancestor of crypts^27^) between cells in different sectors of the colon cause measurable differences in somatic mutation rates.

### Sporadic mutational signatures

Nine signatures (six SBS, one DBS and two ID) were present only in a subset of individuals and/or a subset of crypts and are termed “sporadic”. All were novel, except for SBS2, SBS13 and DBS8. SBS2 and SBS13 are predominantly characterised by C>T and C>G mutations at T<u>C</u>N, are likely due to activity of APOBEC cytidine deaminases and usually occur together^31,32^. SBS2 and SBS13 were clearly observed in one colonic crypt from one individual and one ileal crypt from another, but smaller contributions may be present in additional crypts (Extended Data Fig. 9). To our knowledge, this is the first reported evidence that APOBEC DNA-editing of the human genome occurs in normal cells *in vivo*. However, in the colon at least, it is restricted to a small subset of cells. The factors that initiate it are unknown, although viral entry, retrotransposon transposition and local inflammation have been proposed in other contexts^33^. The wider sequence context of these mutations in normal colon suggests that APOBEC3A is the major contributing enzyme^34^.

Four SBS signatures that do not match the reference set, SBSA-D, were found in normal colorectal cells (SBSA has recently been reported in an oral squamous carcinoma^35^). SBSA is characterised predominantly by T>C mutations at A<u>T</u>A, A<u>T</u>T, and T<u>T</u>T, and T>G mutations at T<u>T</u>T. Its mutation burden correlated closely with that of IDA, which is characterised by single T deletions in short runs of Ts (with a mode of four), suggesting that they are due to the same underlying mutational process. SBSA exhibited a highly variable mutational burden, being present in 29/42 individuals studied, often in just a subset of crypts, and showed evidence of spatial clustering in the colon, with crypts from the same biopsy carrying the signature even though the mutations themselves were not shared (Supplementary Results, Extended Data Fig. 9). 2.5-fold more T>C mutations occurred when the T was on the transcribed than on the untranscribed strand. Transcriptional strand bias is often due to transcription coupled nucleotide excision repair (TC-NER) acting on DNA damaged by exogenous exposures causing covalently bound bulky adducts, but can also be caused by transcription coupled DNA damage^36^. Assuming either is the case, damage to adenine underlies SBSA. To investigate the timing of SBSA mutation generation, phylogenetic trees of mutations were constructed and the mutational signatures in each branch established (Fig. 3, Extended Data Fig. 6). SBSA was confined to early branches of these phylogenies (when these were available for analysis). (Fig. 3b, Extended Data Fig. 6 f, h, z, aa, am, ao). Using the number of SBS1 mutations as indicators of real time, the mutational process underlying SBSA appears to be active usually before 10 years of age. The initiating event for this relatively frequent mutational process is unknown, but the results suggest an extrinsic, locally acting and patchily distributed mutagenic insult occurring during childhood.

**Figure 3.**
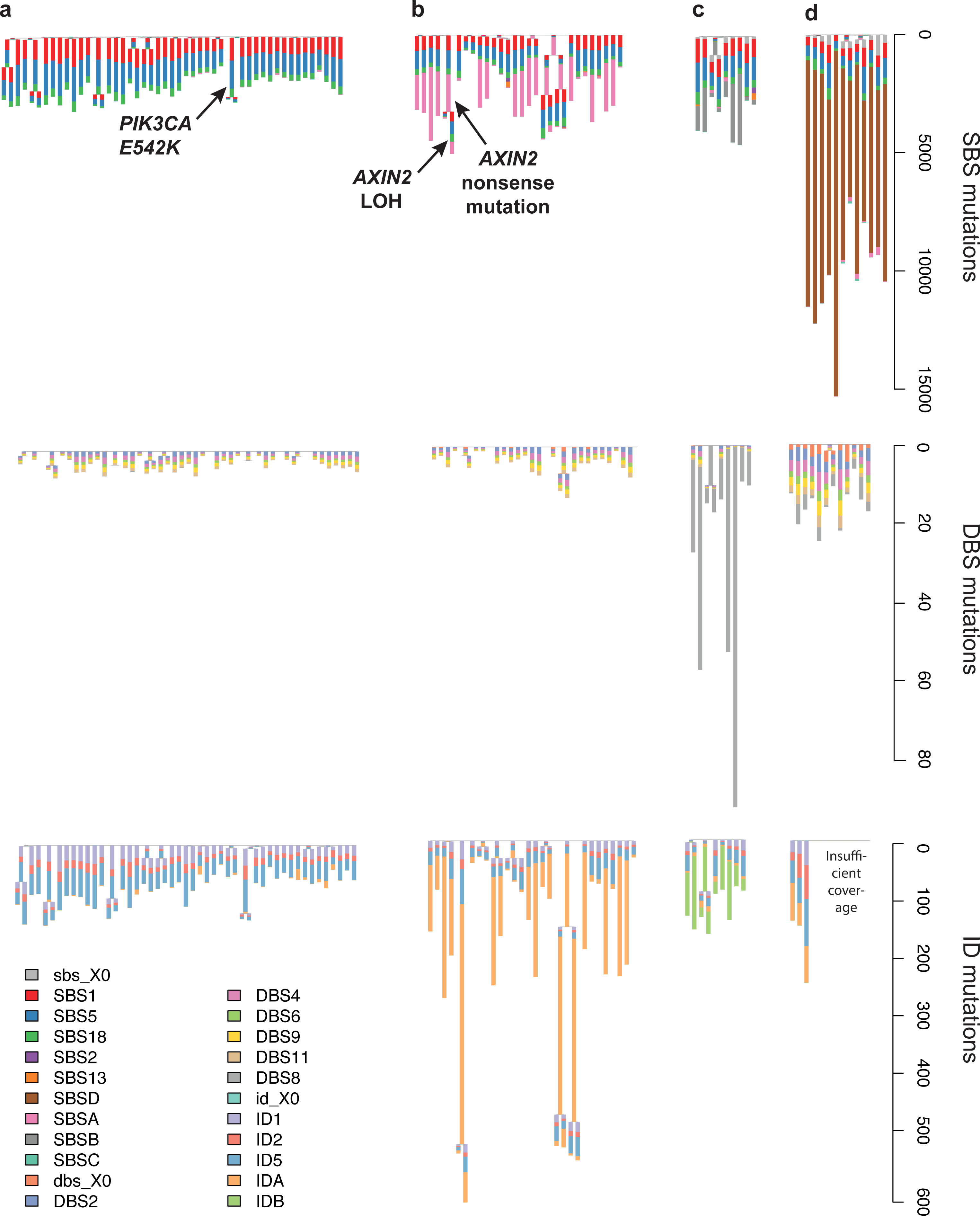
Crypt phylogenies. For four selected individuals (**a–d**), the phylogeny is shown three times: on top, with branch lengths proportional to the number of single base substitutions; in the middle, with branch lengths proportional to the number of doublet base substitutions; on the bottom, with branch lengths proportional to the number of small insertions and deletions. Scale bars are shown on the right–hand side. A stacked barplot of the mutational signatures that contribute to each branch is superimposed onto every branch. Please note that the ordering of signatures along a given branch is just for visualisation purposes: we cannot distinguish the timing of different signatures along a branch. “X0” indicates mutations that could not confidently be assigned to any signature. The phylogenies for all individuals are shown in Extended Data Fig. 6. (**a**) a phylogeny dominated by ubiquitous and known signatures. A *PIK3CA* mutation is shared by two crypts. (**b**) a phylogeny with a strong contribution of SBSA and IDA, as well as an *AXIN2* mutation (the same as in Fig. 4). (**c**) a phylogeny with SBSB, DBS8, and IDB. (**d**) the phylogeny of the individual exposed to chemotherapy, showing a strong contribution of SBSD.

SBSB was predominantly characterised by C>T substitutions at A<u>C</u>A, T>A at C<u>T</u>N, and T>G at G<u>T</u>G and was present in subsets of crypts from four individuals (Fig. 3c, Extended Data Fig. 6). In the two individuals in whom it could be timed (Extended Data Fig. 6 aa, ai), it appeared – as with SBSA – to be active in the first decade of life. SBSB correlated with DBS8 and IDB (Fig. 3c, Extended Data Fig. 9), suggesting that they are caused by the same underlying mutational process. DBS8 is composed of AC>CA and AC>CT mutations and has previously been reported in rare hypermutated cancers with no obvious cause^1^. IDB is dominated by deletion of a single T with no other Ts surrounding it. The mutational process underlying this signature is unknown.

SBSC is characterised predominantly by one C>T mutation in CC dinucleotides. It is of unknown aetiology and primarily affects three crypts from the left colon of one individual with an unremarkable history (Extended Data Fig. 9).

All crypts from a 66 year–old man carried many thousands of mutations of SBSD, characterised predominantly by T>A substitutions with a transcriptional strand bias compatible with damage to adenine. This individual had been treated with multiple chemotherapeutic agents (including cyclophosphamide, doxorubicin, vincristine, prednisolone, chlorambucil, bleomycin and etoposide) for lymphoma and subsequently developed caecal adenocarcinoma. SBSD resembles SBS25, (cosine similarity 0.9), previously found in Hodgkin lymphoma cell lines from two chemotherapy–treated patients^31,37^. To our knowledge this is the first time that the mutational consequences of chemotherapy have been demonstrated in normal human cells *in vivo*. The mutation burden in this individual’s colorectal epithelium was 3–5 fold higher than expected for his age, thus by extrapolation equivalent to that of a 200–300 year–old, and it is plausible that other tissues have been similarly affected.

### Copy number changes and structural variants

Copy number changes and/or structural variants were found in 80 out of 449 (18%) evaluable normal crypts. Five crypts exhibited eight whole chromosome copy number increases which, notably, affected the same three chromosomes: 3, 7 and 9, as well as the X chromosome (Extended Data Fig. 7a). Thus, copy number increases clustered in certain crypts and tended to affect certain chromosomes. No whole chromosome losses were observed. Regions of copy number neutral loss of heterozygosity were observed in 12 crypts, affecting chromosomes 1p, 6p, 7p, 8q, 9q, 10q (twice), 17p, 17q, 18q, 21q and 22q. Five of these copy number changes could be timed and all were estimated to have occurred in adulthood. Two changes that affected the same crypt appeared to be synchronous (Supplementary Results, Extended Data Fig. 7b). Forty–eight large deletions, 18 tandem duplications, four translocations, and two inversions were detected. All were private to a single crypt, except for one deletion which was present in two adjacent crypts sharing few mutations, indicating that the deletion occurred during gestation or early childhood.

### Driver mutations

Driver mutations are those that confer a selective advantage during cancer evolution and may, but need not, promote neoplasia^38^. To search for driver mutations in normal colon, the whole genome sequences of 571 crypts were supplemented with targeted sequencing of 90 known colorectal cancer genes in an additional series of crypts. In total, substitutions in these genes were evaluable in 1,403 crypts and indels in 1,046. Statistical analysis revealed evidence of positive selection on the recessive cancer genes *AXIN2* (three truncating mutations, adjusted q value 0.004) and *STAG2* (two truncating mutations, adjusted q value 0.038) indicating that these mutations are likely drivers. Additional likely driver mutations were identified in cancer genes characterised by canonical missense hotspot mutations. Nine hotspot mutations in *PIK3CA* (E542K, R38H), *ERBB2* (R678Q, V842I, T862A), *ERBB3* (R475W, R667L), and *FBXW7* (R505C, R658Q) were observed (Extended Data Fig. 8). Given the specificity of these hotspot mutations, most are likely to be drivers. In addition, heterozygous truncating mutations were found in the recessive cancer genes *ARID2, ATM* (two), *ATR, BRCA2, CDK12* (two), *CDKN1B, RNF43* (two), *TBL1XR1*, and *TP53* (Supplementary Table). There was no statistical evidence for selection of truncating mutations in the set of 90 colorectal cancer genes overall. The possibility that some have conferred clonal growth advantage, however, is not excluded. No crypt carried more than one putative driver mutation.

23 pairs of adjacent crypts shared over 100 SBS1 mutations and thus were likely to have been generated by postnatal crypt fission. Two pairs carried driver mutations (one with an *AXIN2* nonsense mutation and one with *PIK3CA* E542K), although the association of driver mutations with crypt fission is not significant (p=0.17). In one sister crypt the *AXIN2* mutation was rendered homozygous by copy number neutral chromosome 17q LOH, revealing ongoing clonal evolution in normal colon (Fig. 4, Fig. 3b).

**Figure 4.**
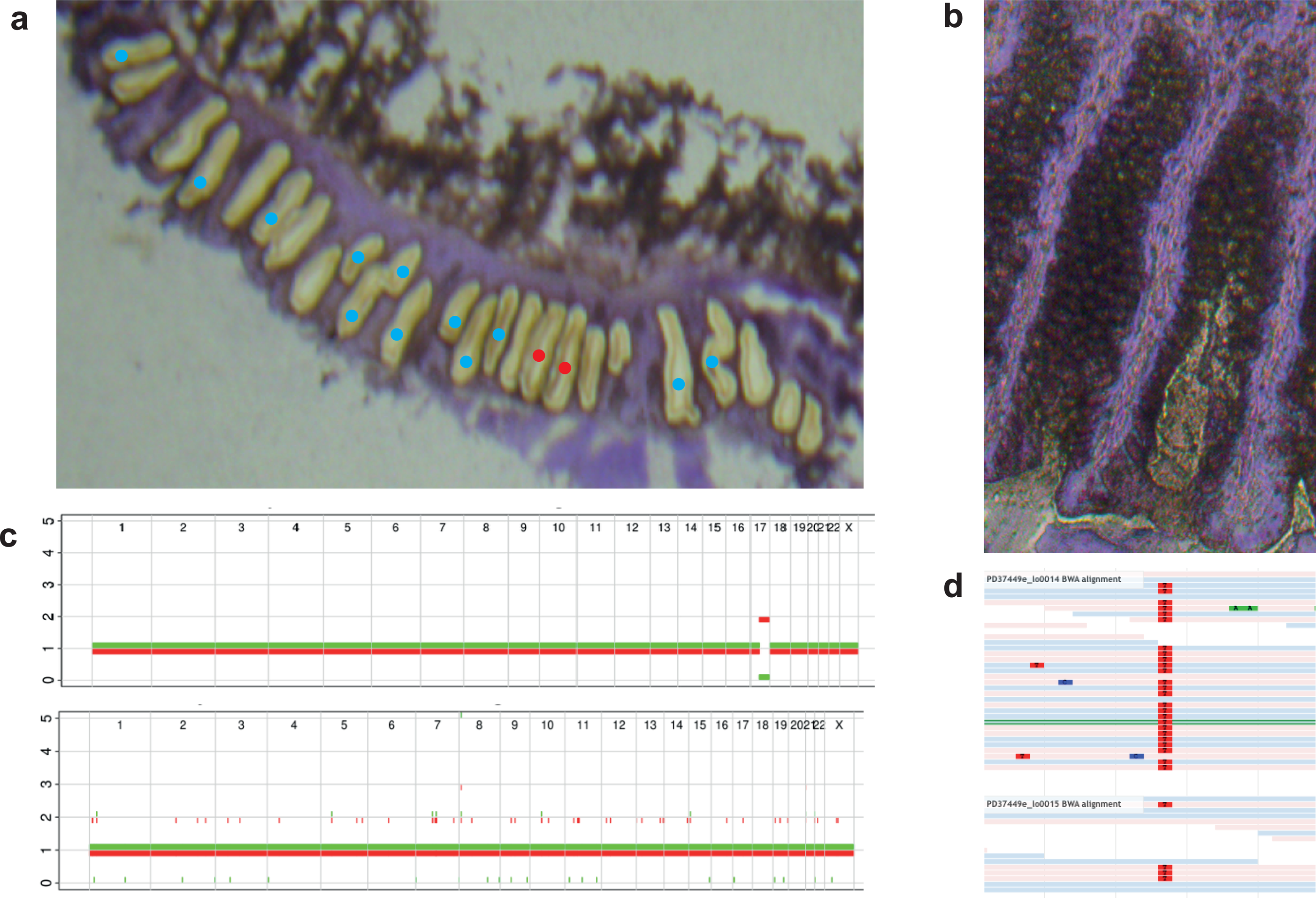
An *AXIN2* driver mutation in normal colon. (**a**) a section (after dissection) in which an inactivating *AXIN2* mutation was found. Red dots represent crypts with the *AXIN2* mutation. Blue dots represent crypts that could be assessed and were found not to have the mutation. Crypts without dots failed sequencing and could not be assessed. (**b**) the two crypts with the *AXIN2* mutations prior to dissection did not appear different to other crypts. (**c**) copy neutral loss of heterozygosity (CNN–LOH) of one of the crypts over the *AXIN2* locus. The copy number state (y axis) for every chromosome is shown, with one allele coloured red and the other green. (**d**) Jbrowse image of reads supporting the *AXIN2* mutations in each of the crypts. The mutation is coloured red. 25 out of 29 reads support the mutation in the crypt that has CNN–LOH; the four reads that do not are presumably the result of stromal contamination.

**Figure 5.**
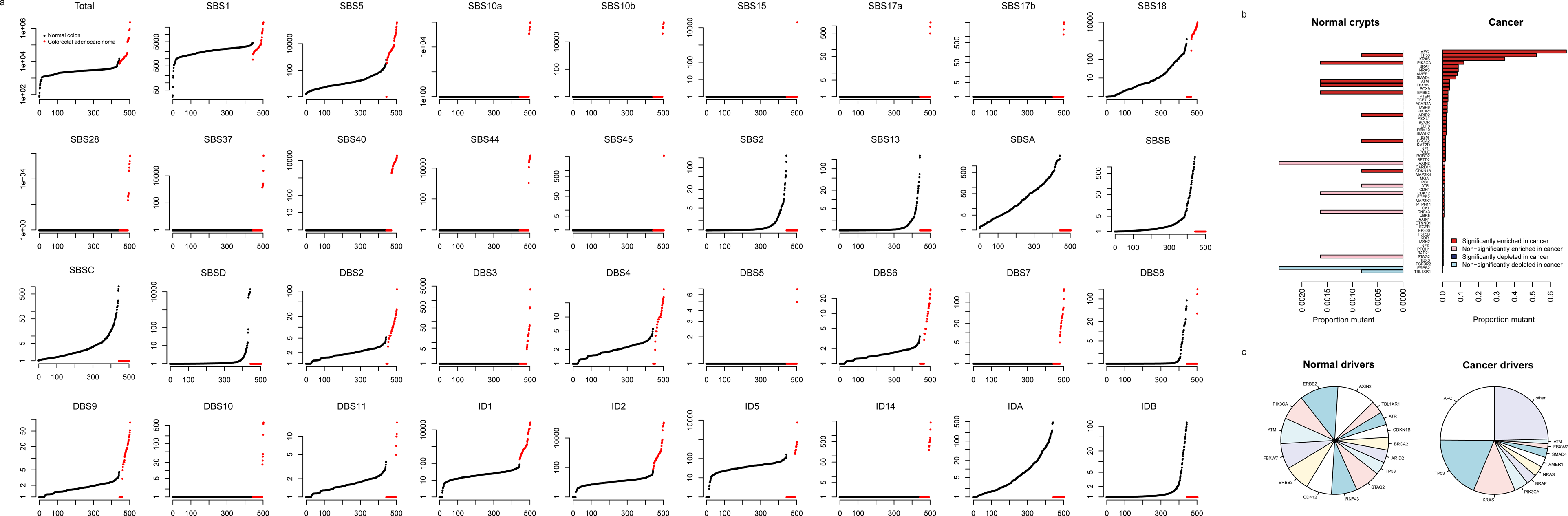
Comparison of the mutational signatures and driver landscape of normal crypts and colorectal adenocarcinomas. (**a**) a comparison of the burden of mutations due to every mutational signature found in either group. For each signature, the (mutation burden+1) of every sample is shown on the y axis on a log scale. Normal colon and cancer samples are ordered within their groups. Colorectal adenocarcinoma signature attributions and burden are from Alexandrov et al.^1^. (**b**–**c**) the frequency of driver mutations in normal colon and colorectal cancer. The frequency of driver mutations is derived using data from The Cancer Genome Atlas Network^43^ (Supplementary Methods). (**b**) the proportion of crypts or cancers with driver mutations in each gene found in either of the two groups. (**c**) the proportion of driver mutations in each gene in normal and cancer.

On the conservative assumption that just the *AXIN2* and *STAG2* truncating mutations and the missense hotspot mutations in *PIK3CA, ERBB2, ERBB3* and *FBXW7* are drivers, ∼1% of normal colorectal crypts (∼150,000 crypts) in a 50–60 year old carries a driver mutation. Since in the over 70s ∼40% of people have an adenoma on colonoscopy^39^ and ∼5% of people develop colorectal cancer over their lifetime^40^ (and some of these may arise from more recently–acquired driver mutations) only an extremely small proportion of these crypt microneoplasms becomes a macroscopically detectable adenoma (< 1/375,000) or carcinoma (< 1/3,000,000) within the following few decades.

The proportion of normal colorectal cells with a driver mutation (1%) is considerably lower than that observed in normal skin (30%). The lower frequency of drivers in colon may be due, at least in part, to the modular structure of glandular epithelia. The small number of stem cells within a crypt diminishes the probability that a cell with a driver mutation will outcompete its wild–type neighbours. Moreover, even if it does colonise the crypt, a mutant stem cell is entombed in it unless it can overcome the largely unknown forces that govern clonal expansion through crypt fission.

### Comparisons with colorectal cancer

There are marked differences between the genomes of normal colorectal stem cells and those of colorectal cancers. The total mutation burdens of base substitutions (10,000–20,000) and indels (1,000–2,000) found in most colorectal carcinomas^1^ (excluding those with hypermutator phenotypes in which it is usually >10–fold more) is higher than the ∼3,000 substitutions and 300 indels found in most normal crypts from 50–60 year old individuals. The particularly high base substitution and indel mutation burdens and associated mutational signatures of DNA mismatch repair deficiency and/or polymerase epsilon/delta mutations were not found in any normal colorectal crypts but are present in ∼20% colorectal cancers. Equally striking is the difference between the 0–4 structural changes per normal crypt (with the majority having none) and the 10s to 100s per colorectal cancer^41^. In all these respects, the genomes of normal crypts with driver mutations were similar to those of normal crypts without drivers (Extended Data Fig. 9).

Elevated mutation burdens are, therefore, characteristic of the evolutionary trajectory from normal colorectal cell to cancer cell. The increased base substitution and indel mutation loads in cancers are due to a combination of higher burdens of the ubiquitous mutational signatures found in normal crypts, additional base substitution and indel signatures thus far found exclusively in cancers (confirming previous reports^5,42^) and larger numbers of copy number changes and structural variation. The causes of some of these additional mutational loads are known (for example, defective DNA mismatch repair and polymerase epsilon/delta mutations) but the remainder are uncertain.

The relative frequencies of mutated cancer genes differ between colorectal adenomas/carcinomas and normal colorectal cells (p=0.003, Supplementary Results). In colorectal cancer, mutations in *APC, KRAS* and *TP53* are common^43^, accounting for 56% of base substitution and indel drivers (Supplementary Methods) but are comparatively rare among normal crypts with driver mutations (1/14). By contrast, mutations in, for example, *ERBB2* and *ERBB3* are relatively common in normal crypts with drivers (5/14) but rare in colorectal cancer (7/631). The results suggest that mutations in *APC, KRAS* and *TP53* confer higher likelihoods of conversion to adenoma and carcinoma than mutations in *ERBB2* and *ERBB3* whereas mutations in *ERBB2* and *ERBB3* confer higher likelihoods of stem cells colonising crypts than *APC, KRAS* and *TP53*. Nevertheless, previous reports suggest that 1:3,500 epithelial cells, and therefore >4,000 crypts per colon, bear *KRAS* G12D^44^, and so even these have a low probability of progression.

## Discussion

This study has characterised all classes of somatic mutation in normal colorectal epithelial stem cells. A substantial repertoire of base substitution and indel mutational processes is operative, some ubiquitous and some sporadic, together with relatively infrequent copy number changes and genome rearrangements. APOBEC DNA–editing occurs in normal colon, albeit only in rare cells. Many signatures, however, are of unknown aetiology and some appear to be acquired early in life. The presence of five times the age–standard mutation load in all colorectal cells, and potentially many other tissues, in an individual who had undergone chemotherapy provides new insight into the impact of such exposures and raises questions pertaining to its relationship with chemotherapy’s relatively modest impact on cancer risk^45^.

The earliest stages of colorectal cancer development have been revealed in this manuscript. They are characterised by numerous crypts carrying driver mutations, of which only a very small fraction ever manifest as macroscopic neoplasms. Certain mutated cancer genes appear to foster this pervasive and invisible wave of microneoplastic change whereas others particularly engender progression to colorectal adenoma and cancer. The conversion of these early microneoplasms to more advanced stages of colorectal neoplasia is associated with acquisition of elevated mutational loads, whether composed of base substitutions, indels, structural variants or copy number changes. More extensive studies of normal colorectal epithelium will enable characterisation of the rarer intermediate stages between these early clones and small adenomas, and refine understanding of the development of the subset of microneoplasms with higher likelihoods of becoming adenomas and carcinomas.

The proportion of normal colorectal epithelial cells with driver mutations is, however, substantially lower than that of other normal tissues so far studied, notably skin^10^ and endometrium^11^. Colorectal epithelium is constituted of crypts, modular units which may themselves constrain clonal expansion, and this architecture may contribute to such differences with skin. The reason for the difference with endometrium, which is also glandular, remains to be explored.

Fundamental questions are being addressed with respect to differences in cancer incidence rates between tissues. The somatic mutation burden in colon and ileum is similar despite the substantially higher cancer incidence rate in colon (as previously noted^4^) and therefore does not appear to account for this difference. Whether the total burden of microneoplastic change across the colon and in other tissues more closely correlates with these differences is yet to be determined.

Finally, this study provides a reference perspective on the mutational signatures and driver mutations in normal colon against which disease states of inflammatory, genetic, neoplastic, degenerative and other aetiologies can be compared. Similar surveys conducted across the range of normal cell types will inform on the universal process of somatic evolution in the human body in health and disease.

## ACKNOWLEDGEMENTS

This work was supported by the Wellcome Trust. We thank Paul Scott, Jo Fowler, David Fernandez–Antoran, and Yvette Hooks for their advice with histology and laser capture microdissection, and Moritz Gerstung for his advice on statistics. We further thank Krishnaa Mahbubani, Rogier ten Hoopen, Cinzia Scarpini, and the Phoenix study team of Nicola Grehan, Irene Debiram–Beecham, Jason Crawte, Tara Nuchkeddy Grant, Pierre Lao–Sirieix, and Andy Hindmarsh for their help with sample collection. Access to transplant organ donor samples were provided by the Cambridge Biorepository for Translational Medicine. Finally, we thank all the individuals who contributed samples to this study.

## AUTHOR CONTRIBUTIONS

MRS and HLS designed the study and wrote the manuscript with contributions from all the authors. KSP, NC, MZ, RCF, NG, FT, AN, MG, and LM recruited patients and obtained samples. PE, RO, HLS, and LM devised the protocol to laser capture microdissect and sequence colonic crypts. HLS prepared sections, microdissected, and lysed colonic crypts. PR contributed to laser capture microdissection. PE and CA made libraries. HLS performed most of the data curation and statistical analysis. MAS devised filters for substitution calling. JW performed in–house NMF signature extraction. TC contributed to statistical analyses. LON provided technical assistance. PJC and IM oversaw statistical analyses. MRS supervised the study.

## EXTENDED FIGURE LEGENDS

**Extended Data Figure 1.**
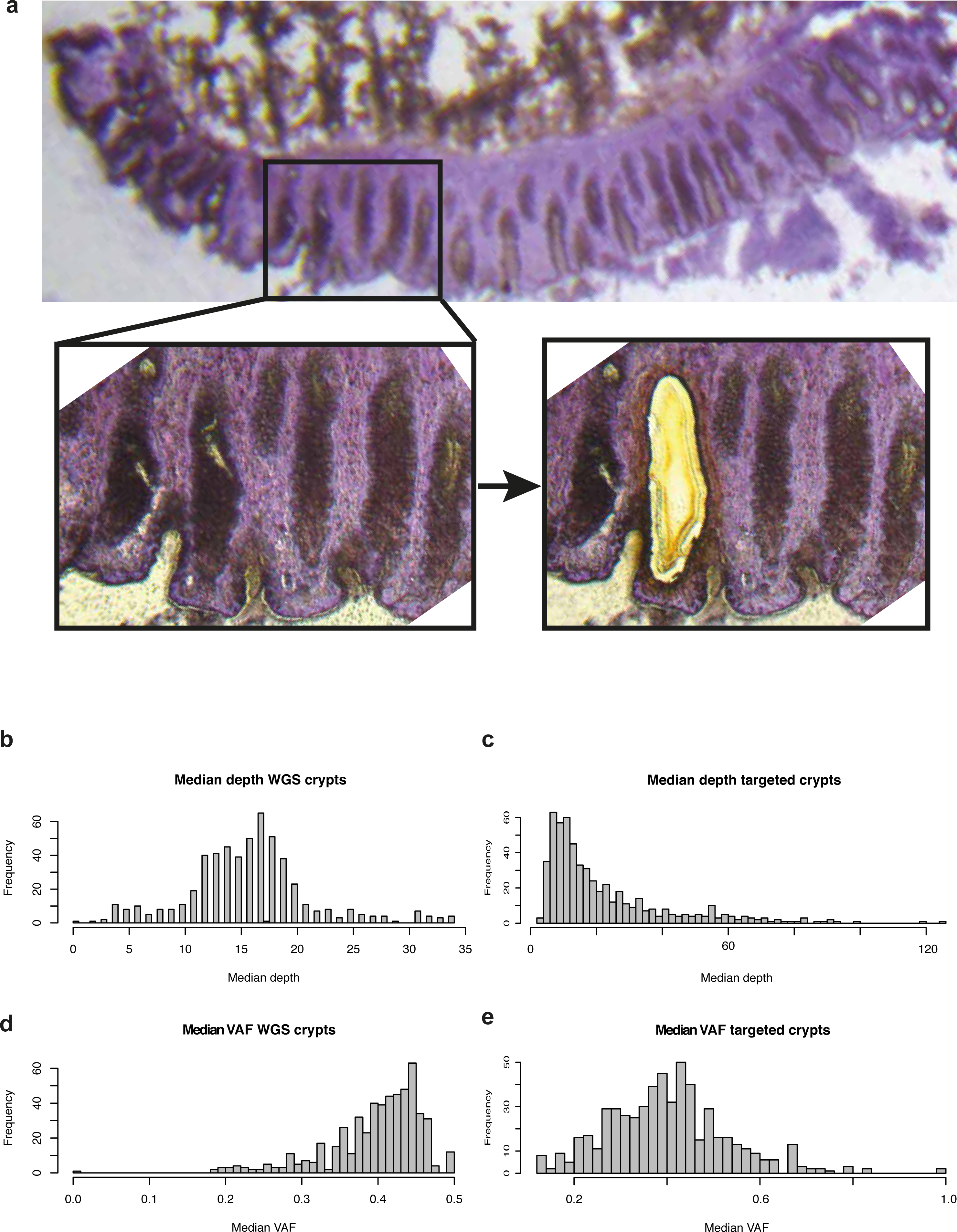
Laser capture microdissection of crypts. **(a)** a representative image of a section of colonic tissue, with a magnified inset showing the section before and after dissection of a crypt. (**b–c**), the coverage of crypts that underwent whole genome **(b)** and targeted **(c)** sequencing. (**d–e**), their respective VAF (which is half of the clonal fraction).

**Extended Data Figure 2.**
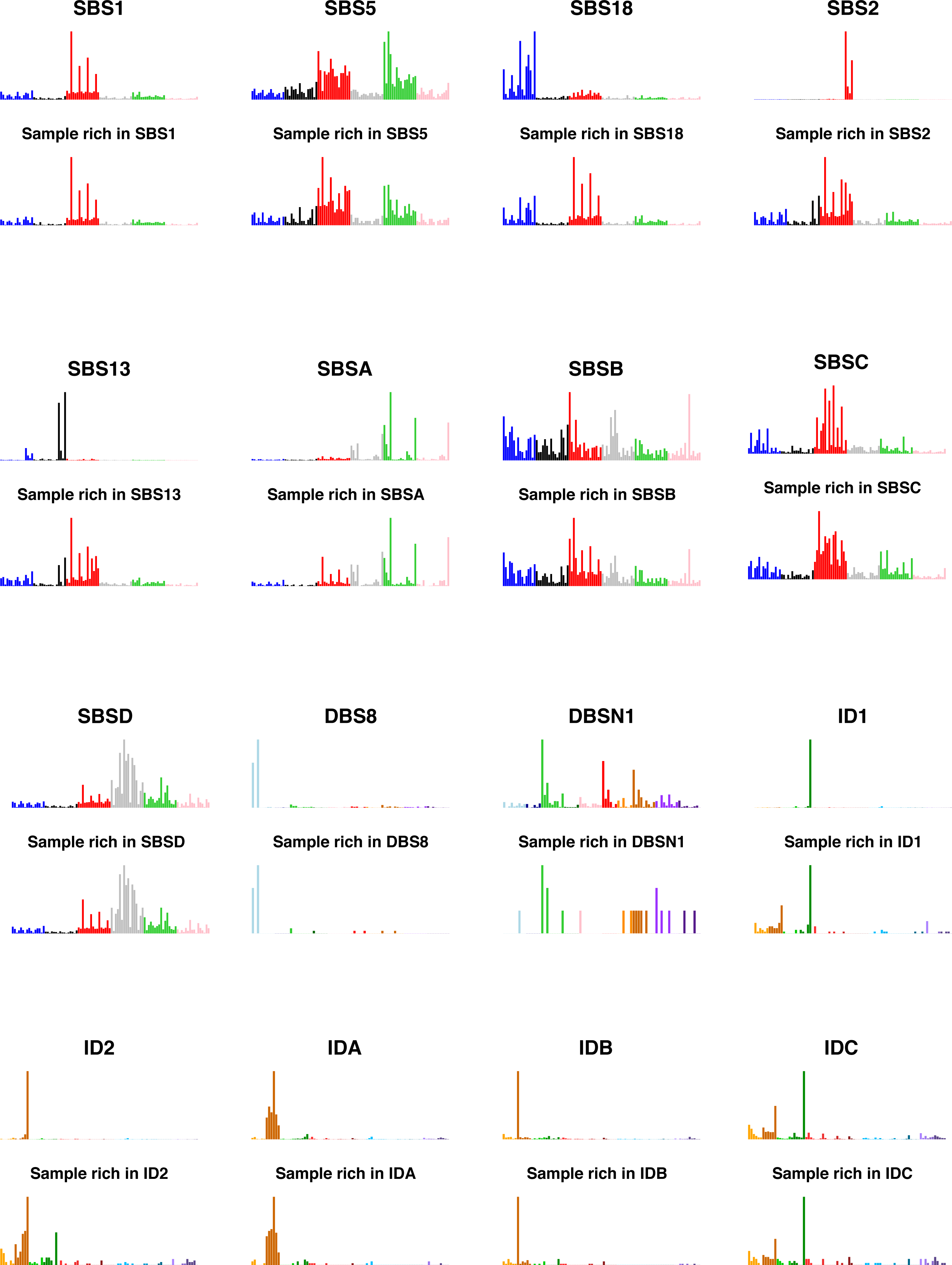
HDP signature extraction results. Results of signature extraction using an HDP with conditioning on signatures known to be active in colorectal cancer. For each signature, the extracted signature and the profile of a sample that has a strong contribution of that signature are shown. Signatures are presented as in Fig. 2. The HDP extraction was followed by deconvolution by Expectation Maximisation (Methods,Extended Data Fig. 3) to produce the version of signatures presented in the main text. HDP, Hierarchical Dirichlet Process.

**Extended Data Figure 3.**
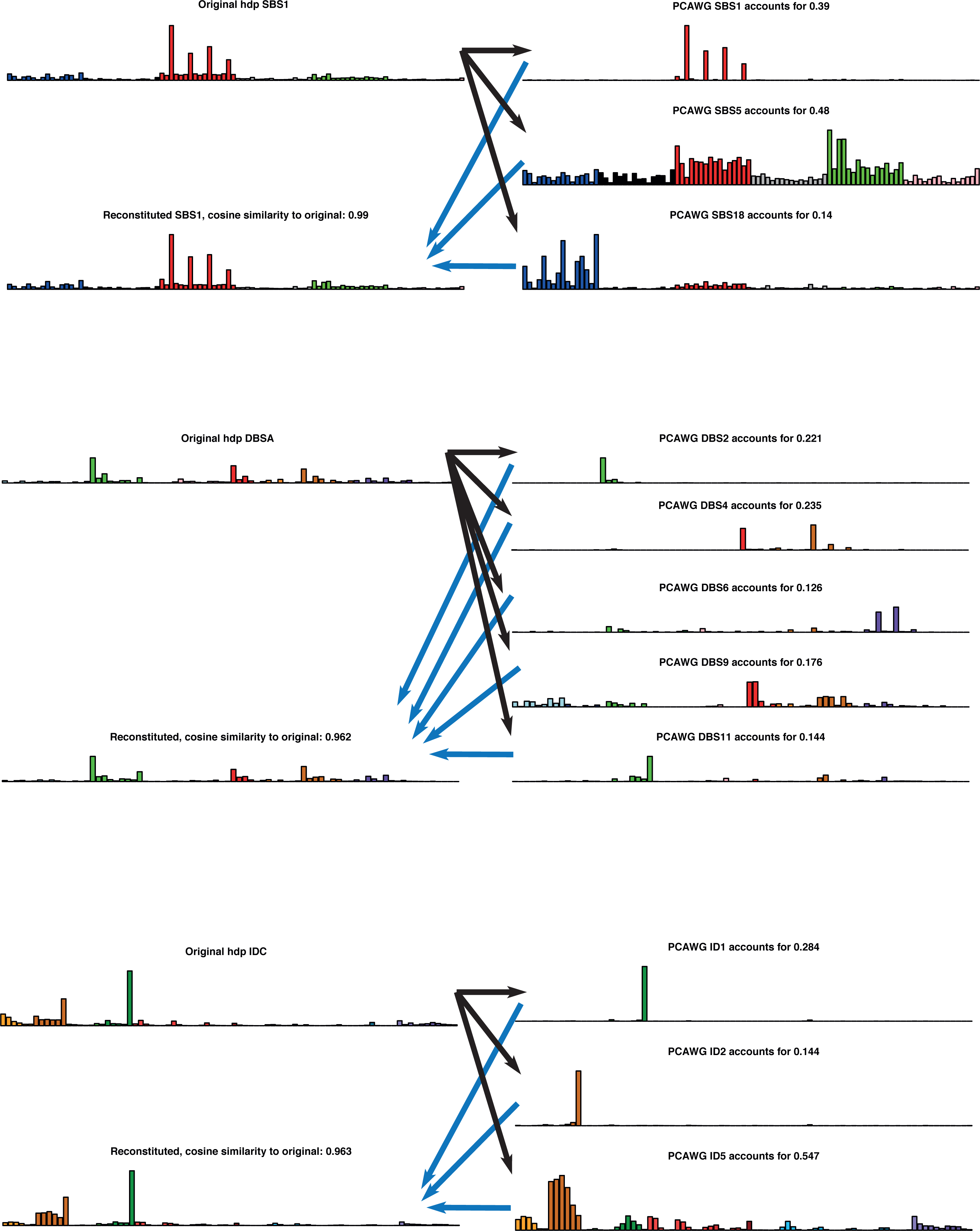
Expectation maximisation decomposition of HDP signatures. Three signatures were decomposed. For each panel, the original HDP version in shown on the top left, the PCAWG signatures that are deemed to contribute at least 10% of mutations to it on the right, and the reconstituted signature built by combining the PCAWG signatures on the bottom left. The cosine similarity of the reconstituted signature to the original is shown in the title to the reconstituted signature plot. HDP, Hierarchical Dirichlet Process; PCAWG, Pan Cancer Analysis of Whole Genomes.

**Extended Data Figure 4.**
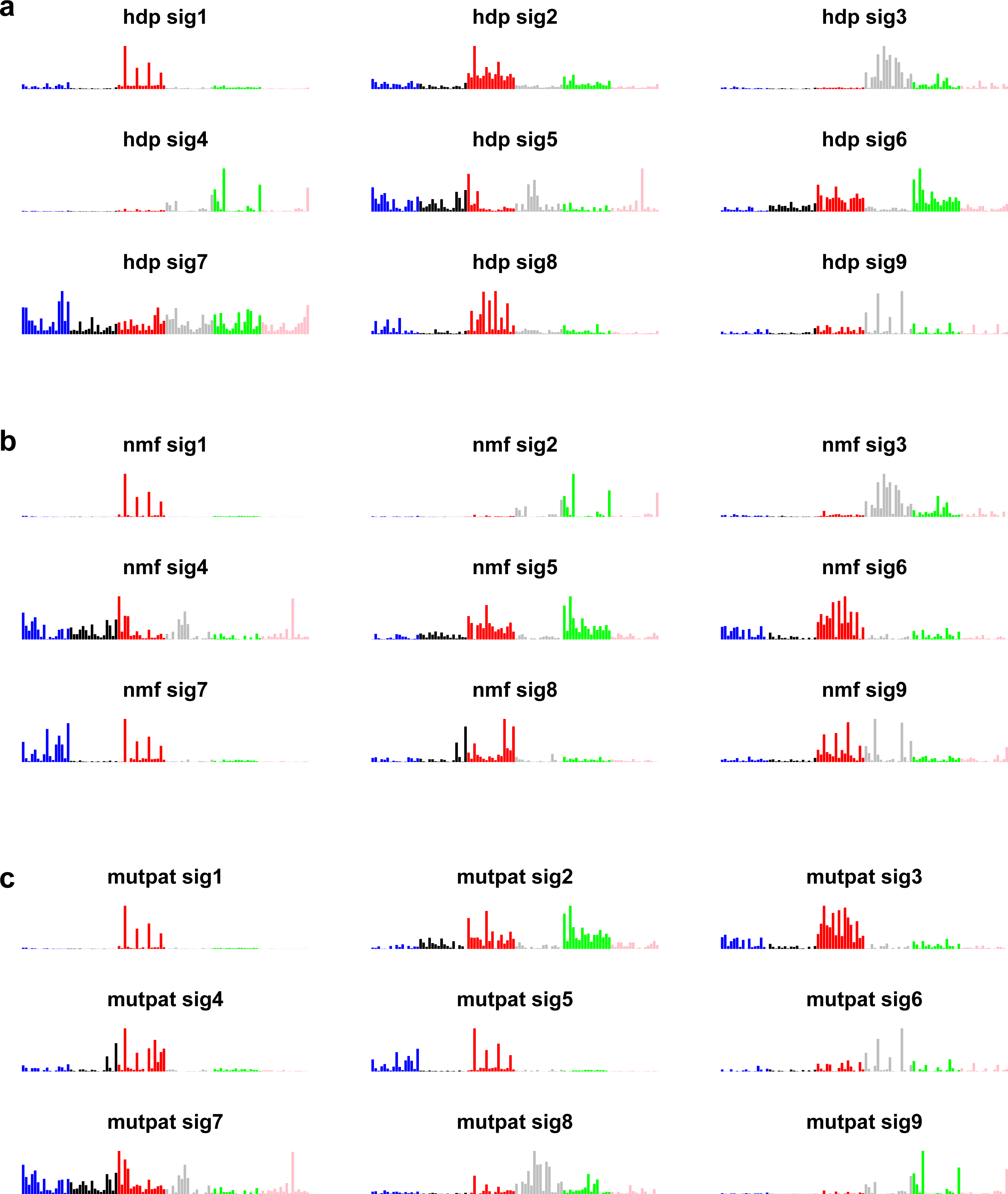
Validation of single base substitution signatures. Other methods of signature extraction were run to test the robustness of signature decomposition. **a,** HDP without pre–conditioning on PCAWG. **b,** In–house NNMF without pre–conditioning on PCAWG. **c**, NNMF implemented by the MutationalPatterns R package (Methods). HDP, Hierarchical Dirichlet Process; PCAWG, Pan Cancer Analysis of Whole Genomes; NNMF, Non–Negative Matrix Factorisation.

**Extended Data Figure 5.**
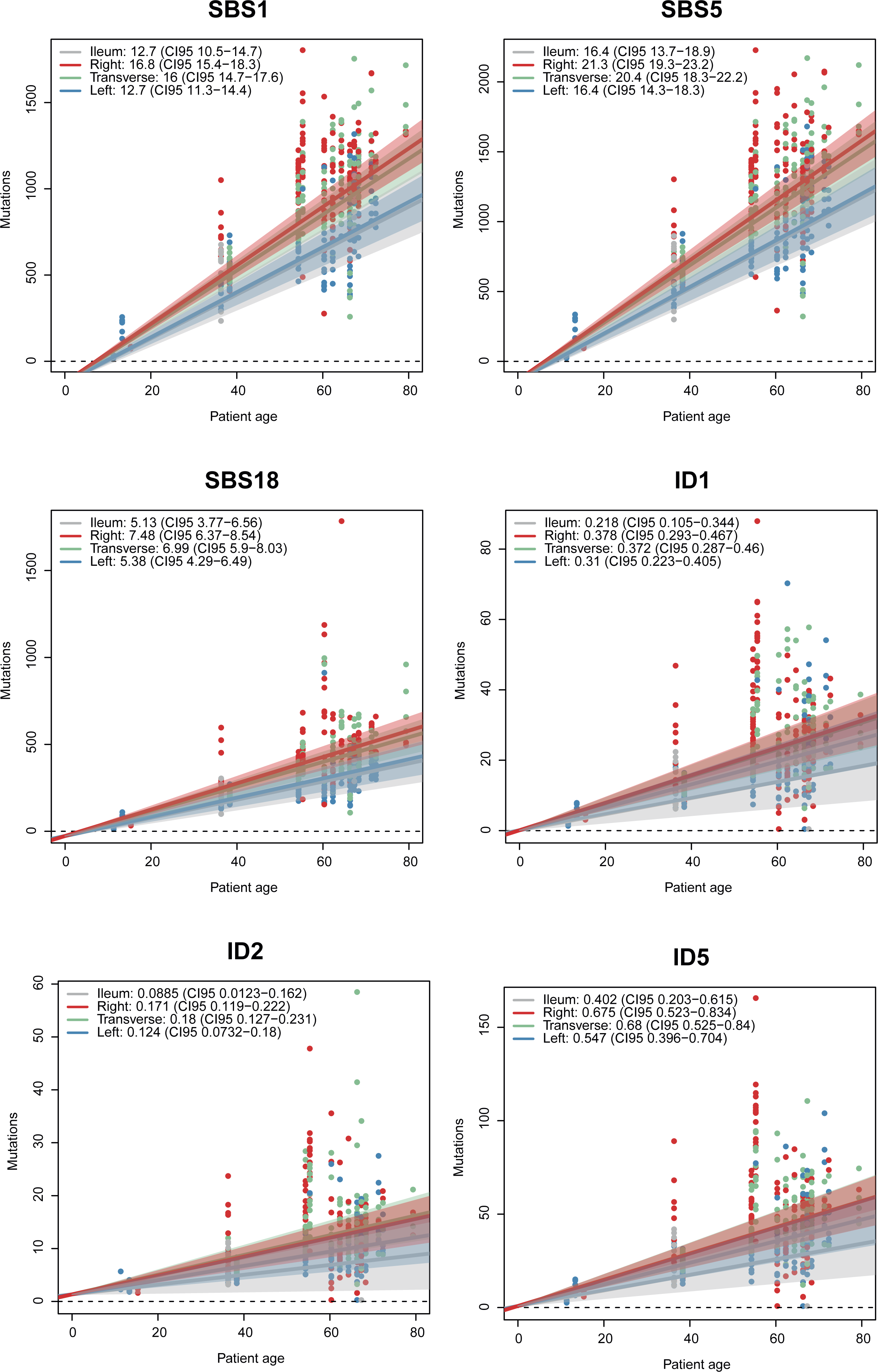
Linear modelling of signature accumulation. For signatures that appeared to show a linear accumulation with age, the mutation rate per site was determined using mixed models, with age and site as fixed effects, and individual as a random effect. Confidence intervals were determined by bootstrapping.

**Extended Data Figure 6.**
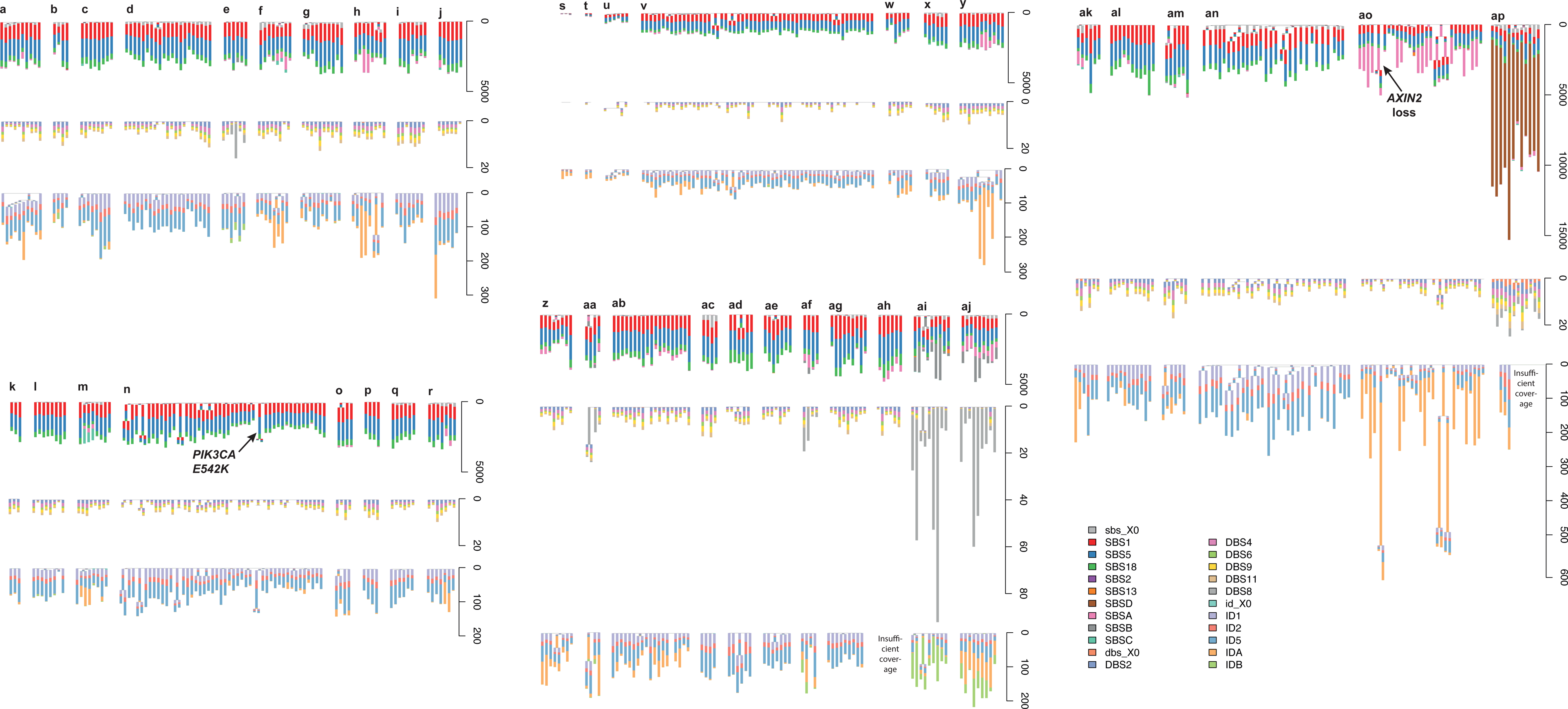
Crypt phylogenies. For every individual, the phylogeny of crypts is shown three times: on top, with branch lengths proportional to the number of single base substitutions; in the middle, with branch lengths proportional to the number of doublet base substitutions; on the bottom, with branch lengths proportional to the number of small insertions and deletions. Scale bars are shown on the right–hand side. A stacked barplot of the mutational signatures that contribute to each branch is overlaid over every branch. “X0” indicates mutations that could not confidently be assigned to any signature. Please note that the ordering of signatures along a given branch is just for visualisation purposes: we cannot distinguish the timing of different signatures along a branch.

**Extended Data Figure 7.**
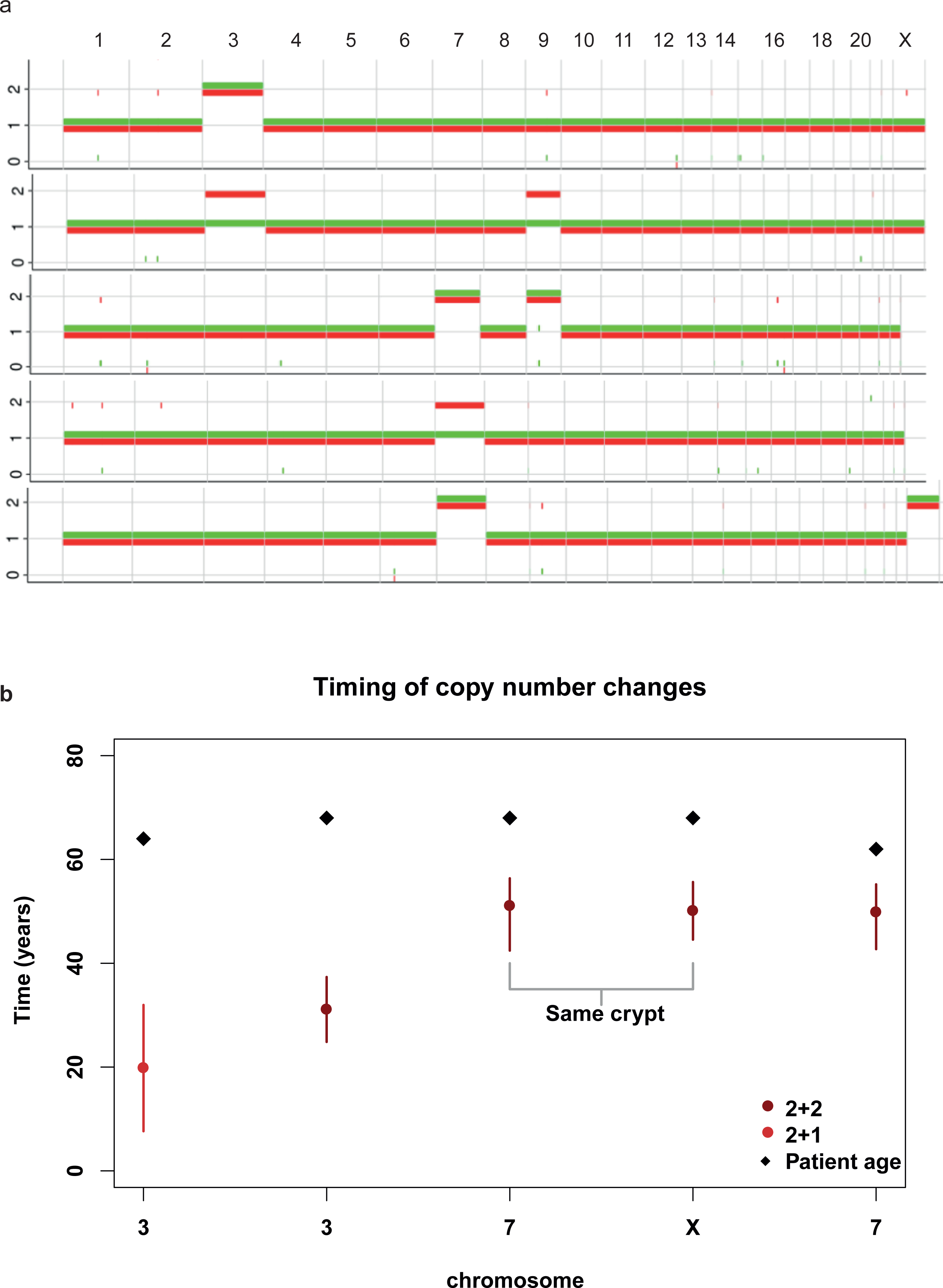
Copy number changes in normal colon. **(a)** whole chromosome amplifications in five crypts. The copy number state (y axis) for each allele, one coloured red, and one coloured green, is shown. Chromosomes are labelled along the top of the graph. **(b)** timing of copy number changes throughout life. Vertical bars represent 95% confidence intervals determined by bootstrapping.

**Extended Data Figure 8.**
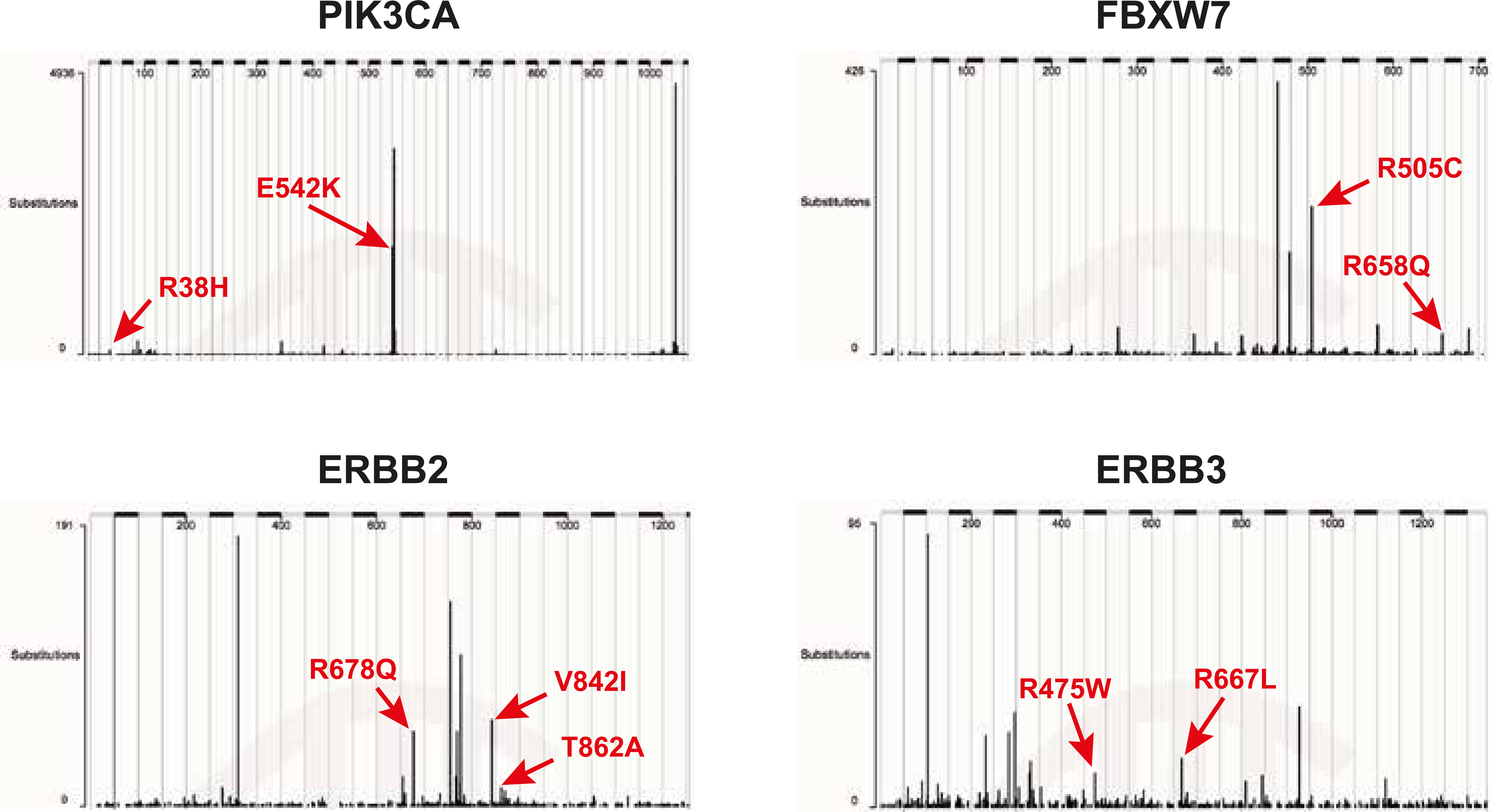
Gain of function driver mutations in normal colon. Putative driver missense mutations in oncogene hotspots. The number of substitutions catalogued in COSMIC are shown on the y axis at each position along the gene, with the mutations observed in our cohort highlighted.

**Extended Data Figure 9.**
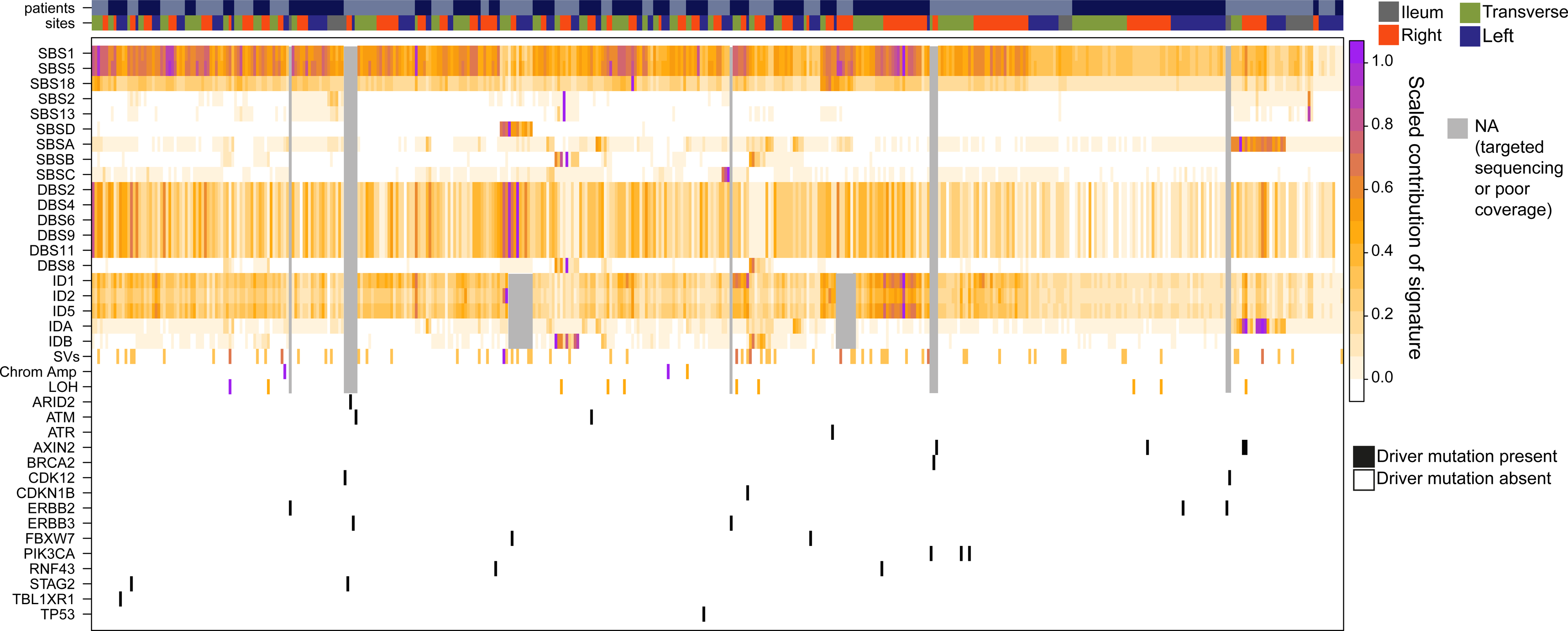
Occurrence matrix of signatures and driver mutations in crypts. For all crypts that were whole genome sequenced to sufficient depth and for crypts that underwent targeted sequencing and in which driver mutations were found, the signatures and driver mutations are shown. Each vertical column represents a crypt. The individual to which each crypt belongs is indicated by alternating colours in the top bar. The site to which each crypt belongs is shown underneath. The contribution of each signature to each crypt;thus the crypt with the largest contribution of a given signature is coloured purple, and the crypt with the smallest contribution is coloured white. Crypts in which the signatures could not be assessed, either because they underwent targeted sequencing or the coverage was poor, are coloured grey. Driver mutations, including heterozygous mutations in tumour suppressor genes, are indicated by a black bar.

## SUPPLEMENTARY METHODS

### Human tissues

We obtained healthy colonic biopsies from four cohorts. The first represents seven deceased organ donors ranging in age from 36 to 67, from whom colonic and small intestinal biopsies were taken at the time of organ donation (REC 15/EE/0152). The second represents individuals aged 60 to 72 who were having a colonoscopy following a positive faecal occult blood test as part of the Bowel Cancer Screening Programme (Ethical approval 08–H0308– 13); we selected 16 who were not found to have either an adenoma or a carcinoma on colonoscopy, and 15 who were found to have a colorectal carcinoma (the normal biopsies that we use were distant from these lesions). The third cohort represents three paediatric patients who underwent routine colonoscopy to exclude inflammatory bowel disease and who were found to have a completely normal intestinal mucosa macroscopically and histologically (REC 12/EE/0482). The final cohort included one 78 year–old gentleman with oesophageal cancer who underwent a warm autopsy (REC 13/EE/0043). All samples were obtained with informed consent and studies approved by East of England Research Ethics Committees.

### Laser capture microdissection of colonic crypts

Fresh frozen biopsies were embedded in optimal cutting temperature (OCT) compound. 30 micrometre sections were fixed in methanol for five minutes, washed three times with phosphate–buffered saline, and stained with Gill’s haematoxylin for 20 seconds. Crypts were isolated by laser capture microdissection, and collected in separate wells of a 96–well plate. They were lysed using the Arcturus PicoPure Kit (Applied Biosystems) according to the manufacturer’s instructions. DNA library prep then proceeded without clean–up or quantification.

### Library preparation

Two library preparation methods were used for laser capture microdissected (LCM) material: in initial experiments sonication was used to fragment DNA, and later, an enzymatic fragmentation method was implemented as it could make libraries from even lower input. Comparison of the two methods showed no difference in mutation calls once post–processing filters (described below) had been implemented. All samples in this study were processed using an Agilent Bravo Workstation (Option B; Agilent Technologies).

For sonication libraries, LCM lysate (20 μl) was mixed with 100 μl TE buffer (Ambion; 10 mM Tris–HCl, 1 mM EDTA) and DNA was fragmented using focused acoustics (Covaris LE220; Covaris, Inc.). Fragmented DNA was mixed with 80 μl Ampure XP beads (Beckman Coulter). Following a 5 min binding reaction and magnetic bead separation, genomic DNA was washed twice with 75% ethanol. Beads were resuspended in 20 μl nuclease–free water (Ambion) and processed immediately for DNA library construction. Each sample (20 μl) was mixed with 2.8 μl of NEBNext Ultra II End Prep Reaction Buffer, 1.25 μl of NEBNext Ultra II End Prep Enzyme Mix (New England BioLabs) and incubated on a thermal cycler for 30 min at 20°C then 30 min at 65°C. Following DNA fragmentation and A–tailing, each sample was incubated for 20 min at 20°C with a mixture of 30 μl ligation mix and 1 μl ligation enhancer (New England BioLabs), 0.9 μl nuclease–free water (Ambion) and 0.1 μl duplexed adapters (100 uM; 5’–ACACTCTTTCCCTACACGACGCTCTTCCGATC*T–3’, 5’–phos–GATCGGAAGAGCGGTTCAGCAGGAATGCCGAG–3’). Adapter–ligated libraries were purified using Ampure XP beads by addition of 65 μl Ampure XP solution (Beckman Coulter) and 65 μl TE buffer (Ambion). Following elution and bead separation, DNA libraries (21.5 μl) were amplified by PCR by addition of 25 μl KAPA HiFi HotStart ReadyMix (KAPA Biosystems), 1 μl PE1.0 primer (100 μM; 5’– AATGATACGGCGACCACCGAGATCTACACTCTTTCCCTACACGACGCTCTTCCGA TC*T–3’) and 2.5 μl iPCR–Tag (40 μM; 5’– CAAGCAGAAGACGGCATACGAGATXGAGATCGGTCTCGGCATTCCTGCTGAACC GCTCTTCCGATC–3’) where ‘X’ represents one of 96 unique 8–base indexes The sample was then mixed and thermal cycled as follows: 98 °C for 5 min, then 12 cycles of 98 °C for 30 s, 65°C for 30 s, 72 °C for 1 min and finally 72 °C for 5 min. Amplified libraries were purified using a 0.7:1 volumetric ratio of Ampure Beads (Beckman Coulter) to PCR product and eluted into 25 μl of nuclease–free water (Ambion). DNA libraries were adjusted to 2.4 nM and sequenced on the HiSeq X platform (illumina) according to the manufacturer’s instructions with the exception that we used iPCRtagseq (5’– AAGAGCGGTTCAGCAGGAATGCCGAGACCGATCTC–3’) to read the library index.

For enzymatic fragmentation, LCM lysate (20 ul) was mixed with 50 ul Ampure XP beads (Beckman Coulter) and 50 μl TE buffer (Ambion; 10 mM Tris–HCl, 1 mM EDTA) at room temperature. Following a 5 min binding reaction and magnetic bead separation, genomic DNA was washed twice with 75% ethanol. Beads were resuspended in 26 μl TE buffer and the bead/genomic DNA slurry was processed immediately for DNA library construction. Each sample (26 μl) was mixed with 7 μl of 5X Ultra II FS buffer, 2 μl of Ultra II FS enzyme (New England BioLabs) and incubated on a thermal cycler for 12 min at 37°C then 30 min at 65°C. Following DNA fragmentation and A–tailing, each sample was incubated for 20 min at 20°C with a mixture of 30 μl ligation mix and 1 μl ligation enhancer (New England BioLabs), 0.9 μl nuclease–free water (Ambion) and 0.1 μl duplexed adapters (100 uM; 5’– ACACTCTTTCCCTACACGACGCTCTTCCGATC*T–3’, 5’–phos–GATCGGAAGAGCGGTTCAGCAGGAATGCCGAG–3’). Adapter–ligated libraries were purified using Ampure XP beads by addition of 65 μl Ampure XP solution (Beckman Coulter) and 65 μl TE buffer (Ambion). Following elution and bead separation, DNA libraries (21.5 μl) were amplified by PCR by addition of 25 μl KAPA HiFi HotStart ReadyMix (KAPA Biosystems), 1 μl PE1.0 primer (100 μM; 5’– AATGATACGGCGACCACCGAGATCTACACTCTTTCCCTACACGACGCTCTTCCGA TC*T–3’) and 2.5 μl iPCR–Tag (40 μM; 5’– CAAGCAGAAGACGGCATACGAGATXGAGATCGGTCTCGGCATTCCTGCTGAACC GCTCTTCCGATC–3’) where ‘X’ represents one of 96 unique 8–base indexes The sample was then mixed and thermal cycled as follows: 98 °C for 5 min, then 12 cycles of 98 °C for 30 s, 65°C for 30 s, 72 °C for 1 min and finally 72 °C for 5 min. Amplified libraries were purified using a 0.7:1 volumetric ratio of Ampure Beads (Beckman Coulter) to PCR product and eluted into 25 μl of nuclease–free water (Ambion). DNA libraries were adjusted to 2.4 nM and sequenced on the HiSeq X platform (Illumina) according to the manufacturer’s instructions with the exception that we used iPCRtagseq (5’– AAGAGCGGTTCAGCAGGAATGCCGAGACCGATCTC–3’) to read the library index.

### Whole genome sequencing

We generated paired end sequencing reads (150bp) using Illumina XTEN^®^ machines resulting in ∼15x coverage per sample. Sequences were aligned to the human reference genome (NCBI build37) using BWA–MEM.

### Targeted sequencing

A 2.3 MB capture panel was designed in–house to pull down genes that are known or suspected to play a role in neoplasia. We performed custom RNA bait design following the manufacturer’s guidelines (SureSelect, Agilent). Samples were multiplexed on flow cells and subjected to paired end sequencing (75–bp reads) using Illumina HiSeq2000 machines. One 96–well plate of samples was sequenced on each lane, but as tissue recovery was variable, a range of coverage was achieved. Sequences were aligned to the human reference genome (NCBI build37) using BWA–align.

### Data Availability

Whole genome and targeted sequencing data are deposited in the European Genome Phenome Archive (EGA). sequencing data have been deposited with EGA accession EGAD00001004192, EGAD00001004192, and EGAD00001004193.

### Code Availability

Code for statistical analyses is provided as part of the supplement. Custom R scripts and their input data for signature analysis are available on GitHub at https://github.com/HLee– Six/colon_microbiopsies. All other code is available from the authors on request.

### Calling substitutions

Substitution calling was broken down into three steps: mutation discovery; filtering to produce a list of clean sites; and genotyping, where the presence or absence of every mutation in every sample is evaluated.

First, mutations were initially discovered using the Cancer Variants through Expectation Maximisation (CaVEMan) algorithm^46^. CaVEMan uses a naïve Bayesian classifier to derive the probability of all possible genotypes at each nucleotide. CaVEMan copy number options were set to major copy number 5 and minor copy number 2 for normal clones, as in our experience this maximises sensitivity. The algorithm was run using an unmatched normal in order to be able to derive phylogenies: had another sample from the same individual been treated as a matched normal, early embryonic mutations would have been treated as germline and discarded, resulting in incorrect trees.

Second, a number of post–processing filters were applied. These included filtering against a panel of 75 unmatched normal samples to remove common single nucleotide polymorphisms, post–processing as described previously^32^ and two filters (only applied to whole genome sequencing data) designed to remove mapping artefacts associated with BWA–MEM: the median alignment score of reads supporting a mutation should be greater than or equal to 140, and fewer than half of these reads should be clipped. The library preparation protocol for microbiopsies produced shorter library insert sizes than standard methods. Reads could therefore overlap, resulting in double counting of mutant reads. Fragment–based statistics were generated to prevent the calling of variant supported by a low number of fragments. Variants were annotated by ANNOVAR^47^ and fragment–based statistics (fragment coverage, number of fragments supporting the variant, fragment–based allele fraction) were calculated for each variant after the exclusion of marked PCR duplicates. In the rare event of discordance in the called base at the variant position between overlapping paired–end reads,the base with the highest quality score was selected. Fragment–based statistics were calculated separately for high quality fragments (alignment score ≥ 40 and base scores ≥ 30). Variants supported by at least three high quality fragments were retained and used for the next stage of variant filtering. Inspection of variants specific to LCM experiments revealed that the vast majority were present within inverted repeats capable of forming hairpin structures, that they were supported by reads with very similar alignment start position (and so not marked as PCR duplicates), and were primarily located close to the alignment start within the supporting reads. Commonly these variants coincided with other proximal variants (1–30 bp), but filtering based on variant proximity would also remove actual kataegis events. *In silico* modelling of the potential hairpin showed that the variants were aligning to each other in the stem of the structure, but could not form a base pair, while all other bases could. The artefacts are likely the consequence of erroneous processing of cruciform DNA (existing either prior to DNA isolation or formed during library preparation) by the enzymatic digestion protocol applied. We have considered modelling the hairpin structures to filter these variants, but given the fact that read clustering (i.e., similar alignment position) serves as a hallmark for these artefacts, we opted to use the proximity of the variant to the alignment start, and the standard deviation (SD) and median absolute deviation (MAD) of the variant position within the supporting reads, as features for filtering. These statistics were calculated separately for positive and negative strand aligned reads. In case the variant was supported by a low number of reads (i.e., 0–1 reads) for one of the strands, the filtering was based only on the statistics generated for the other strand. Per variant, if one of the strands had too few reads supporting, it was required for the other strand that either: (I) there should be ≤ 90% supporting reads to report the variant within the first 15% of the read starting from the alignment start, or (II) the statistics MAD > 0 and SD > 4. Per variant, if both strands were supported by sufficient reads it was required for both strands separately that either: (I) there should be ≤ 90% supporting reads to report the variant within the first 15% of the read, (II) the statistics MAD > 2 and a SD > 2, or (III) that the other strand should have the statistics MAD > 1 and SD > 10 (i.e., the variant is retained if the other strand demonstrates strong measures of variance). In our experience, the proposed strategy vastly reduces the number of artefactual variants while retaining all other variants, as assessed by running the last filtering step on WGS data from non–LCM experiments.

Third, mutations were genotyped in every sample. A pileup of all the samples from a given individual was constructed, counting the number of mutant and wild type reads in every sample over every site that had been called in any sample from that person. Only reads with a mapping quality of 30 or above and bases with a base quality of 30 or above were counted. After applying these filters, mutations were genotyped based on the number of mutant and wild type reads at each locus. Mutations were called based on a variant allele fraction (VAF) > 0.2, a depth > 7, and at least 4 mutant reads. If the depth over a locus was less than seven in a given sample, or if there was more than one mutant read but the other criteria were not met, the genotype was set to NA for tree construction purposes. Loci that were set to NA in more than one third of the samples were removed for construction of the phylogeny. Positions were called as germline if they were either called as present or NA in all of the samples from a given individual.

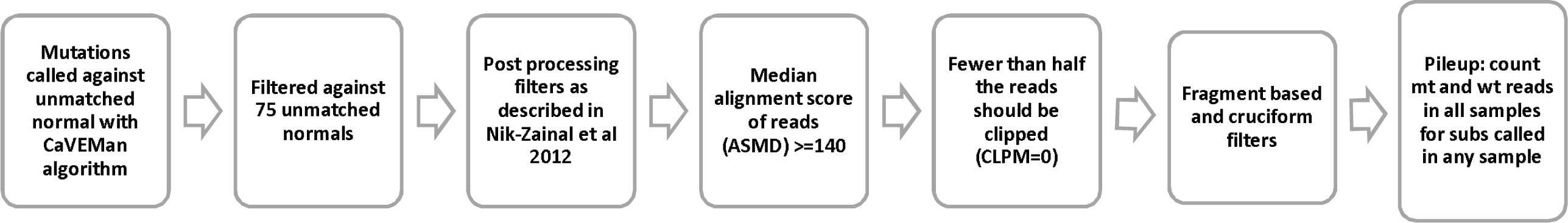

### Calling short insertions and deletions (indels)

As for substitutions, calling of indels was broken down into mutation discovery, filtering, and genotyping. Mutations were called with the Pindel algorithm^48^ using an unmatched normal. Post processing filters were applied as in Nik–Zainal et al.^32^, and the number of mutant and wild–type reads was tabulated as above. The same dataset–specific filters were applied as for substitutions. Indels were then genotyped based on a VAF>0.2, a depth of at least 10, and support of at least 5 mutant reads.

### Calling structural variants

Genomic rearrangements were called using the BRASS algorithm^41^ (https://github.com/cancerit/BRASS). Abnormally paired read pairs from WGS were grouped and filtered by read remapping. Read pair clusters with ≥50% of the reads mapping to microbial sequences were removed, as were rearrangements where the breakpoint could not be reassembled. Candidate breakpoints were matched to copy number breakpoints defined by ASCAT (see below) within 10kb. Only structural variants where the two breakpoints were more than 1000 base pairs apart were considered. Structural variants were called against a matched normal skin or blood sample when available and against another crypt from the same individual with good coverage when not.

### Calling copy number

Copy number changes were called using the Allele–Specific Copy number Analysis of Tumours (ASCAT) algorithm^49^. The same matched normal sample was used as for calling structural variants. For additional validation of copy number changes in normal colon, the QDNAseq algorithm^50^ was run. ASCAT uses both the read depth and ratios of heterozygous single nucleotide polymorphisms to determine an allele–specific copy number, while the QDNAseq relies solely on variations in sequencing coverage. To call amplifications and deletions in the colonic microbiopsy cohort, only those that were both called by ASCAT and showed a clear departure from the background log2ratio by QDNAseq were retained. To call copy neutral loss of heterozygosity in this cohort, all such events called by ASCAT were checked visually on Jbrowse^51^ to verify an imbalance of parental snps. Only crypts with >10X coverage, for which copy number changes could be reliably detected, were used.

### Detection of driver variants and positive selection

Driver mutations were detected both through an unbiased dNdS method and through manual annotation. For these analyses, the CaVEMan and Pindel calls were used without post– processing filters in order to maximise our sensitivity. All putative driver variants were visually inspected using Jbrowse^51^, and so we could afford a higher false positive rate in the mutation discovery phase.

dNdScv^52^ was used to conduct three tests: first, using only the whole genome sequencing data, an analysis of selection over all genes; second, using combined whole genome and targeted sequencing data, over all the genes covered by the bait–set; and finally, using again this combined dataset, over 90 selected cancer genes (appendix). R code for this analysis is included in the supplementary information.

Manual annotation of driver variants based on prior knowledge complemented this. A list of 90 colorectal cancer genes (appendix) curated from the literature that were also covered by the bait–set were intersected with the list of substitutions and indels from combined whole genome and targeted sequencing. Mutations were annotated as putative drivers if they were either missense mutations that fell in an oncogene hotspot (based on visualisation of the distribution of mutations in the gene on COSMIC^53^), or if they were truncating mutations that fell in a tumour suppressor gene.

Structural variants that might act as drivers were assessed by intersection of genes involved in each structural variant with the twelve genes involved in gene fusions that have been reported in colorectal cancer in COSMIC (*VTI1A, TCF7L2, TPM3, NTRK1, PTPRK, RSPO3, ETV6, NTRK3, EIF3E, RSPO2, C2orf44*, and *ALK*). No fusion genes were found. None of the genes involved in structural variants in our data overlapped with the list of 90 cancer genes used for assessing substitutions and indels, and nor were there any genes that were affected by more than one structural variant. No high–level copy number amplifications were observed and there were no homozygous deletions.

### Estimation of frequency of driver mutations in cancer

Publically–available colorectal cancer mutation calls were obtained from The Cancer Atlas Network^43^. Driver mutations were annotated manually in the same way as in our dataset: only mutations that fell in the 90 genes that we had selected were considered, and they were annotated as putative drivers if they were either missense mutations that fell in an oncogene hotspot (based on visualisation of the distribution of mutations in the gene on COSMIC^53^), or if they were truncating mutations that fell in a tumour suppressor gene.

### Construction of phylogenies

Phylogenies are used in this analysis for timing mutations. The most informative branches in this case are the long branches shared by a small number of crypts, which are very robust to all tree construction methods. Trees were built using maximum parsimony using substitutions called as described above. For every individual, the input matrix of mutation calls was bootstrapped 100 times. Phylogenies were constructed for each replicate using the Wagner method of the Mix programme from the Phylip suite of tools^54^. The consensus of all the phylogenies constructed was used.

The phylogenies were validated using the indel calls. To do this, the same procedure as for substitutions was followed for indel matrices. As there were fewer indels than substitutions, nodes in indel phylogenies were generally reconstructed with lower confidence than in substitution phylogenies, but they broadly agree. 85% of nodes reconstructed with >=90% confidence in the indel tree were present with exactly the same set of descendants in the substitution trees.

The phylogeny inference programme used provided the topology of the tree but not the assignment of mutations. Mutations from the input matrix of genotypes therefore have to be re–assigned to branches. In order to assign a set of mutation calls with no false negative and no false positives to a tree, each branch of the tree was considered in turn. If a mutation was called in all the descendants of a given branch, and in no samples that were not descendants of the branch, mutations were assigned to that branch.

Some colonic microbiopsies suffered from low coverage and stromal contamination. For this reason, we did not expect mutations to fit the tree perfectly, as a mutation that was truly present in a colony might be missed if too few supporting reads are found. Mutations were only assigned to the tree in order to determine the mutational processes active at a particular time. We reasoned that it was preferable to assign only mutations that fit the tree perfectly and adjust the branch lengths based on the power to call mutations at a given branch, rather than attempting to assign mutations that fit the tree imperfectly. Using the clonality and coverage of all descendants of a branch, the proportion of true substitutions or indels on the branch that would be first discovered (whether by CaVEMan or Pindel) and then genotyped as present according to the criteria described above was calculated. The observed branch length was then adjusted by dividing by this proportion. This was done for both substitutions and indels, but not for structural variants and for larger copy number changes due to a lack of data: most branches have no large variants and so could not be extended appropriately. Rearrangements and copy number changes were assigned to phylogenies manually.

### Extraction of mutational signatures

Mutational signatures were extracted using the mutations assigned to every branch of a phylogeny as a ‘sample’. This allows better discrimination of mutational processes that may occur at different times within the same cell. Mutations were categorised following the method used by the Mutational Signatures working group of the Pan Cancer Analysis of Whole Genomes (PCAWG)^1^. Single base substitutions were categorised into 96 classes according the identity of the pyrimidine mutated base pair, and the base 5’ and 3’ to it. Doublet base substitutions were categorised into 78 classes according to the identity of the reference and alternative bases. Indels were classified according to whether they were an insertion or a deletion, the identity of the inserted/deleted base, the length of the mononucleotide tract in which they occurred, or the degree of homology with the surrounding sequence into 83 classes (Fig. 1a).

Signatures were extracted using a hierarchical Dirichlet Process^55,56^. Code and the input mutations are provided at https://github.com/HLee–Six/colon_microbiopsies. First, the algorithm was conditioned on the set of mutational signatures that have found to be operative in colorectal cancers in PCAWG^1^: SBS1, SBS2, SBS3, SBS5, SBS13, SBS16, SBS17a, SBS17b, SBS18, SBS25 (included although it is not found in colorectal cancer because the similarity with the mutational profile with crypts from one individual had been previously noted), SBS28, SBS30, SBS37, SBS40, SBS41, SBS43, SBS45, SBS49, DBS, DBS3, DBS4, DBS6, DBS7, DBS8, DBS9, DBS10, DBS11, ID1, ID2, ID3, ID4, ID5, ID6, ID7, ID8, ID10,and ID14. This allows simultaneous discovery of new signatures and matching to known ones. Nine single base substitution (SBS), two doublet base substitution (DBS), and five indel (ID) signatures were discovered (Extended Data Fig. 2). Despite pre–conditioning, signatures that were perfectly correlated in all samples were still amalgamated. This occurred, for example, with signatures 1, 5, and 18. Therefore, expectation maximisation was used to deconvolute all HDP signatures into known PCAWG signatures. If a signature reconstituted from the components that expectation maximisation extracted (only including PCAWG signatures that accounted for at least 10% of mutations in each sample to avoid over–fitting) had a cosine similarity to the HDP signature of more than 0.95, the signature was presented as its expectation maximisation deconvolution. Three HDP signatures met these criteria: the HDP SBS1 signature was deconvoluted into a mixture of PCAWG SBS1, PCAWG SBS5, and PCAWG SBS18; the HDP DBSA was deconvoluted in PCAWG DBS2, PCAWG DBS4, PCAWG DBS6, PCAWG DBS9, and PCAWG DBS11; and the HDP IDC was deconvoluted into PCAWG ID1, PCAWG ID2, and PCAWG ID5 (Extended Data Fig. 3). To test the robustness of this signature analysis, other signature extraction methods were used: HDP with no pre–conditioning, the non–negative matrix factorisation (NNMF) method used by Blokzijl and colleagues^4^, and a version of the NNMF algorithm used by Alexandrov and colleagues^1^. These all produced comparable results (Extended Data fig. 4).

### Statistical analyses

All statistical analyses were performed in R (Supplementary Results).

